# TMS-EEG signatures of glutamatergic neurotransmission in human cortex

**DOI:** 10.1101/555920

**Authors:** Franca König, Paolo Belardinelli, Chen Liang, Debora Desideri, Florian Müller-Dahlhaus, Pedro Caldana Gordon, Carl Zipser, Christoph Zrenner, Ulf Ziemann

## Abstract

Neuronal activity in the brain is regulated by an excitation-inhibition balance. Glutamate is the main excitatory neurotransmitter. Transcranial magnetic stimulation (TMS) evoked electroencephalographic (EEG) potentials (TEPs) represent a novel way to quantify pharmacological effects on neuronal activity in the human cortex. Here we tested TEPs under the influence of a single oral dose of two anti-glutamatergic drugs, perampanel, an AMPA-receptor antagonist, and dextromethorphan, an NMDA-receptor antagonist, and nimodipine, an L-type voltage-gated calcium channel blocker in 16 healthy adults in a pseudorandomized, double-blinded, placebo-controlled, crossover design. Single-pulse TMS was delivered to the left motor cortex and TEPs were obtained pre-and post-drug intake. Dextromethorphan specifically increased the amplitude of the N45, a negative potential around 45 ms after the TMS pulse, while perampanel reduced the P70 amplitude in the non-stimulated hemisphere. Nimodipine and placebo had no effect on TEPs. These data extend previous pharmaco-TMS-EEG studies by demonstrating that the N45 is regulated by a balance of GABAAergic inhibition and NMDA-receptor-mediated glutamatergic excitation. In contrast, AMPA-receptor-mediated glutamatergic neurotransmission contributes to interhemispherically propagated activity reflected in the P70. These data are important to understand the physiology of TEPs as markers of excitability and propagated activity in the human cortex in health and disease.

## Introduction

Transcranial magnetic stimulation (TMS) evoked electroencephalographic (EEG) potentials (TEPs) reflect excitability and effective connectivity of the human brain (Ilmoniemi and Kicic 2010; Rogasch and Fitzgerald 2013; Chung et al. 2015; Tremblay et al. 2019). However, the exact physiological mechanisms underlying the multiple TEPs evoked by, e.g., motor cortex stimulation (Bonato et al. 2006; Lioumis et al. 2009) remain still largely elusive. Pharmaco-TMS-EEG has demonstrated that the N45, a negative potential around 45 ms after the TMS pulse, is regulated by GABAAergic inhibition as its amplitude is enhanced by allosteric positive modulators at GABAA receptors, such as benzodiazepines and zolpidem (Premoli et al. 2014; Premoli et al. 2018), but reduced by the experimental compound S44819 (Darmani et al. 2016), a specific antagonist at the alpha-5 subtype of the GABAA receptor. In contrast, GABABergic inhibition contributes to the N100, as its amplitude at the site of the stimulated motor cortex is increased by baclofen, a specific GABAB receptor agonist (Premoli et al. 2014; Premoli et al. 2018). The P25 seems to reflect corticospinal excitability as its amplitude correlates with the amplitude of motor evoked potentials (MEPs) measured with electromyography (EMG) (Mäki and Ilmoniemi 2010; Cash et al. 2017), and is suppressed by the voltage-gated sodium channel blocker carbamazepine (Darmani et al. 2018a). Finally, late TEPs, in particular the P180, are also suppressed by voltage-gated sodium channel blockers (Premoli et al. 2017; Darmani et al. 2018a).

The excitatory glutamatergic system has so far not been tested with TMS-EEG, although it plays a fundamental role in the excitation-inhibition balance to regulate neuronal excitability in cerebral cortex (Tatti et al. 2017). Understanding the role that glutamatergic neurotransmission plays on TEP generation is essential to obtain an accurate physiological understanding of the TMS evoked EEG potentials. This is of relevance if TEPs shall be used as diagnostic/prognostic markers in psychiatric or neurological disorders (Tremblay et al. 2019), many of which show a dysfunction in the glutamatergic system, e.g., schizophrenia (Hasan et al. 2014), epilepsy (Eid et al. 2008) or amyotrophic lateral sclerosis (Blasco et al. 2014).

Here, we investigated the effects of a single oral dose of two anti-glutamatergic drugs (perampanel, dextromethorphan) and the L-type voltage-gated calcium channel (L-VGCC) blocker nimodipine (Hess et al. 1984) on TEPs in healthy subjects in a pseudorandomized double-blind placebo-controlled crossover design. Perampanel is a selective, non-competitive postsynaptic α-amino-3-hydroxy-5-methyl-4-isoxazole propionic acid (AMPA) receptor antagonist (Rogawski and Hanada 2013). Dextromethorphan is a prodrug whose active metabolite, dextrorphan, acts as a non-competitive N-methyl-D-aspartate (NMDA) receptor antagonist (Wong et al. 1988). AMPA and NMDA receptors are the main ionotropic receptors for glutamate in the central nervous system. AMPA receptor-mediated currents generate fast excitatory postsynaptic potentials (EPSPs), while NMDA receptor activation provides a prolonged EPSP that can last several hundred milliseconds. Action potential generation is largely controlled by AMPA receptor de/activation, while the longer kinetics of NMDA receptors enable spatial and temporal summation of postsynaptic potentials (Niciu et al. 2012). Accordingly, perampanel is used as an antiepileptic drug (Faulkner 2017), while dextromethorphan has demonstrated efficacy in reducing synaptic plasticity in human cortex (Stefan et al. 2002; Wankerl et al. 2010; Weise et al. 2017). Finally, L-VGCCs are not significantly involved in controlling the release of glutamate from presynaptic nerve terminals (Catterall 2011) but block synaptic plasticity in human cortex (Wolters et al. 2003; Wankerl et al. 2010; Weise et al. 2017), probably through inhibition of calcium flux into depolarized postsynaptic cells (Igelmund et al. 1996). We had no specific hypotheses as to the effects of these study drugs on TEP amplitudes, given that pharmaco-TMS-EEG is a nascent field. Therefore, the study is exploratory, but positive findings would significantly enhance our understanding of the mechanisms underlying TEPs, since the effects of anti-glutamatergic drugs and VGCC blockers have not been tested in this context.

## Material und methods

### Participants

Eighteen male participants (mean age ± SD: 26.0 ± 3.5 years, range: 22-36 years), were included in this study. All subjects underwent physical and neurological examination, and were screened for possible contraindications to TMS (Rossi et al. 2009) and to the study medication. Inclusion criteria were written informed consent, right-handedness (mean laterality score ± SD: 88 ± 15 % according to the Edinburgh Inventory (Oldfield 1971)) and male gender, to avoid possible effects of the menstrual cycle on cortical excitability (Smith et al. 1999). Exclusion criteria were: presence or history of neurologic and psychiatric disease, use of illicit or recreational drugs, smoking, and a history of low blood pressure (assessed with history of past measurements or symptoms, e.g. syncope). The study was approved by the Ethics Committee of the Medical Faculty of Eberhard-Karls-University Tübingen (registration number 526/2014BO1). Sixteen subjects completed all the experimental sessions. One participant did not finish the study due to medical conditions unrelated to the study and one other subject dropped out during the measurements. Therefore, the data analyses are based on 16 subjects.

### Experimental design

A combined pharmaco-TMS-EEG approach (Premoli et al. 2014; Darmani et al. 2016) with a pseudorandomized, placebo-controlled, double-blinded crossover design was employed to study the acute effects of perampanel, dextromethorphan and nimodipine on TEP amplitudes.

Each experimental session consisted of one pre-and one post-drug measurement, which involved the same procedures, as follows. Before each measurement, resting motor threshold (RMT), defined as the minimum intensity sufficient to elicit an MEP amplitude ≥ 50 μV in at least five out of ten trials was determined, using the relative frequency method (Groppa et al. 2012). Then, resting-state EEG (3 min eyes open) was recorded, followed by the delivery of 150 single monophasic TMS pulses with a random interstimulus interval of 5 ± 1 s for TEP recordings. The TMS target was the hand area of the left primary motor cortex (M1), a constant coil position was maintained throughout the experiment. The pre- and post-drug measurements were separated by the administration of the study drug, immediately after the pre-drug measurements, and a pause, which allowed the drug to reach peak serum level (see Supplementary Material and Fig. 1). Due to different pharmacokinetics, drugs and/or placebo were applied at two different time points to ensure a double-blinded design.

**Figure 1.**
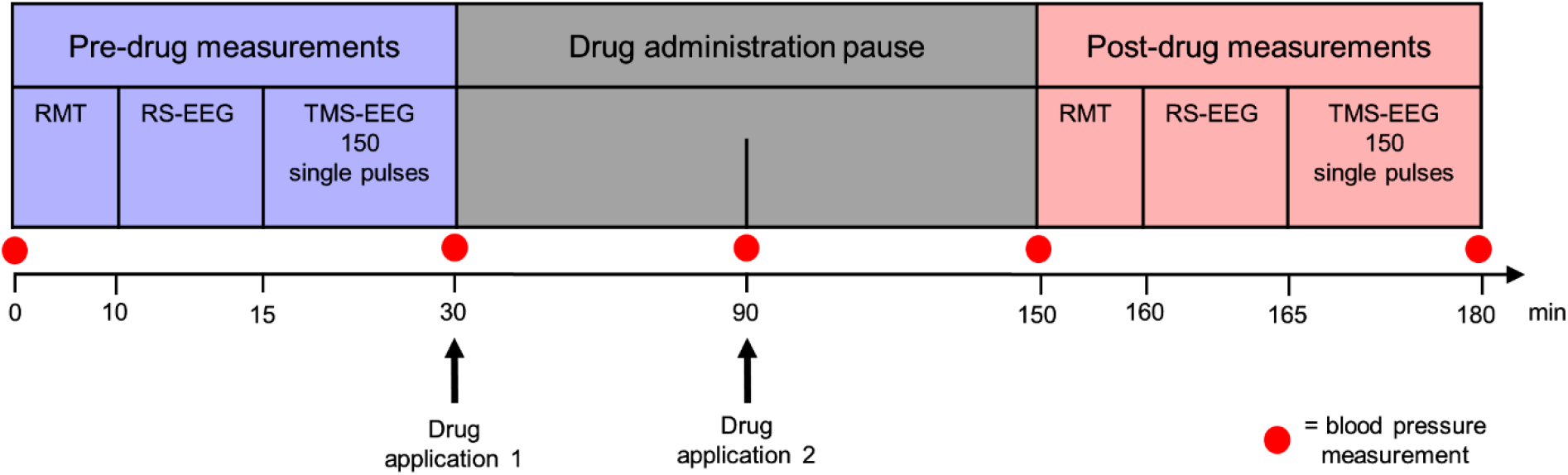
Timeline of experiments. Determination of resting motor threshold (RMT) at the beginning of pre- and post-drug measurements was followed by resting-state EEG (RS-EEG) and a block of 150 single TMS pulses over the left primary motor cortex with simultaneous EEG measures (TMS-EEG). During the two-hour medication pause two drug administrations were performed (see Supplementary Table 2). One hour after the second drug administration, the post-drug measurements were obtained in the same sequence as the pre-drug measurements. Blood pressure was monitored throughout the measurement.

Participants received a single oral dose of perampanel (12 mg/6 mg, Fycompa^®^, Eisai Pharma), dextromethorphan (120 mg, Hustenstiller-ratiopharm^®^ Dextromethorphan, ratiopharm GmbH), nimodipine (30 mg, Nimodipin-Hexal^®^, Hexal AG), or placebo (P-Tabletten Lichtenstein; Placebo Kapseln). Drug dosages employed in the study are approved for medical use. The order of drugs was pseudorandomized and balanced across subjects. Based on drug pharmacokinetics reported in the literature the study drugs are characterized by different peak-plasma times (Supplementary Table 1). Accordingly, study drugs and placebo were given at the two time points indicated in Supplementary Table 2. To avoid carry-over drug effects, consecutive sessions in each participant were separated by at least two weeks.

### TMS-EEG and EMG data recordings

All participants were seated in a comfortable reclining chair throughout pre- and post-drug measurements. They were instructed to keep their eyes open and to focus on a small black cross in front of them to reduce eye movements. Their right hand was comfortably placed and relaxed throughout the experiment to avoid muscle activation, as this increases the MEP amplitude (Hess et al. 1987).

A TMS-compatible EEG amplifier (BrainAmp DC, BrainProducts GmbH, Munich, Germany) and 62 high-density TMS-compatible C-ring slit EEG electrodes (EASYCAP, Germany) arranged in the International 10-20 montage (Dmochowski et al. 2017) were used to acquire EEG at a sampling rate of 5 kHz. To monitor eye movement and blinking, two additional electrodes where placed above the right eye and at its outer canthus. All electrode impedances were maintained at < 5 kΩ throughout the session. In order to avoid possible EEG contamination by auditory evoked potentials caused by the TMS coil discharge click (Nikouline et al. 1999), white noise was delivered to the participants through earphones during the TMS-EEG recordings (Massimini et al. 2005; Casarotto et al. 2010). The sound pressure level was calibrated until participants indicated that they could no longer hear the TMS clicks.

TMS stimuli were applied to the hand knob of the left M1 using a focal figure-of-eight coil (external loop diameter: 90 mm). The coil was connected through a BiStim module with a Magstim 200^2^ magnetic stimulator (all devices from Magstim Co, Whitland, Dyfed, UK) with a monophasic current waveform. The coil was oriented with the handle pointing backwards and 45° away from the midline, to induce current in the brain oriented from lateral-posterior to anterior-medial (Di Lazzaro et al. 2008). The optimal coil position to elicit MEPs in the right abductor pollicis brevis (APB) muscle was determined as the site that produced consistently the largest MEPs using a stimulation intensity slightly above RMT (motor “hotspot”) (Groppa et al. 2012). MEPs were recorded through surface EMG electrodes (Ag-AgCl cup electrodes) in a belly-tendon montage. The EMG signal was recorded using the Spike2 software (Cambridge Electronic Design). The EMG raw signal was amplified (Digitimer D360 8-channel amplifier), bandpass filtered (20 Hz - 2 kHz) and digitized at an A/D rate of 10 kHz (CED Micro 1401; Cambridge Electronic Design). For constant coil placement throughout the experiment, the coil position at the APB hotspot was marked on the EEG cap. All TMS pulses were applied to the APB hotspot at an intensity of 100 % RMT (Premoli et al. 2014; Darmani et al. 2016; Darmani et al. 2018b), to limit possible contamination of TEPs by re-afferent signals from MEPs (Fecchio et al. 2017). The RMT was re-tested at the beginning of the post-drug measurements (Fig. 1) and, if different from pre-drug RMT, TMS intensity was adjusted to keep the re-afferent signals similar across pre- and post-drug measurements. The inter-trial interval was 5 s ± 25 % random variation to limit habituation.

### Data processing

EEG data processing and analysis were performed using customized analysis scripts on MATLAB R2016a and the Fieldtrip open source MATLAB toolbox (Oostenveld et al. 2011). The continuous EEG data was segmented into epochs from −600 to 600 ms relative to the TMS pulse. EEG data from 1 ms before to 15 ms after the TMS pulse were removed and spline interpolated (Thut et al. 2011). Afterwards, data was down-sampled to 1000 Hz. Bad trials and noisy channels were removed by means of visual inspection of the EEG epochs (mean percentage of removed epochs ± SD: 25.4 ± 12.0 %; mean number ± SD of removed channels: 4.5 ± 2.5). Then, independent component analysis (ICA) was applied to the EEG data in a two-steps procedure (Rogasch et al. 2014). In a first ICA step, TMS related artefacts were removed (mean number of removed components ± SD: 4.3 ± 2.6). Subsequently the data was filtered with a 1-80 Hz Butterworth zero phase band pass filter (3^rd^ order) and a 49-51 Hz notch filter. ICA was then performed again and components representing physiological (i.e., eye blinking or eye movements, muscle artifacts), electrical or small amplitude TMS related artefacts were removed (mean number of removed components ± SD: 13.6 ± 6.2). Successively, removed channels were interpolated using the signal of the neighboring channels (Perrin et al. 1989) and data were re-referenced to linked mastoids (average of EEG electrodes TP9 and TP10). Finally, data were baseline-corrected by subtracting the average of the signal in the time window from 600 to 100 ms prior to the TMS pulse (Premoli et al. 2014) and were smoothed with a 45 Hz low pass filter (Butterworth zero phase band pass filter, 3^rd^ order). TEPs were analyzed channel-wise, by averaging the EEG data of all retained trials, separately for the pre- and post-drug measurements.

For MEP analysis, EMG data were epoched from −100 to 100 ms around the TMS pulse. An epoch was discarded if the absolute value of the mean EMG signal 100 to 0 ms before the TMS pulse exceeded a pre-innervation threshold > 0.02 mV. The mean percentage (± SD) of discarded epochs due to pre-innervation was 11.0 ± 17.9%.

### Statistics

Five non-overlapping time windows of interest (TOIs) were *a priori* defined based on the group average TEPs across subjects, pre- and post-drug measurements, the four drug sessions and all EEG channels. TOIs were centered around the latencies of the canonical M1 TEP peaks P25, N45, P70, N100 and P180 (Komssi et al. 2004a; Bonato et al. 2006; Premoli et al. 2014). Specifically, TOIs were set at 16-34 ms (P25), 38-55 ms (N45), 56-82 ms (P70), 89-133 ms (N100), and 173-262 ms (P180) after the TMS pulse. For each condition, drug-induced TEP modulations were evaluated for each individual TOI using channel-wise paired-sample t-tests. Family-wise error rate (FWER) was controlled by using a cluster-based permutation approach (Maris and Oostenveld 2007), as implemented in Fieldtrip. This approach tests the null hypothesis that data in the experimental conditions are drawn from the same probability distribution and clusters the t-values resulting from the paired-sample t-tests that exceed an *a priori* defined threshold of p < 0.05, based on neighboring channels and time points. The minimum number of channels below the significance threshold to form a cluster was 2. The t-statistics at cluster level was then computed summing the t-values within each cluster and comparing the maximum of the obtained t-values. A reference distribution of the maximum of the cluster t-values was obtained by re-applying the same procedure on the data randomized across the pre-drug vs. post-drug measurements. We used 1500 randomizations to obtain the reference distribution and rejected the null hypothesis with p < 0.05 if less than 5 % of the permutations used to construct the reference distribution yielded a maximum cluster-level t-value larger than the one observed in the original data. The same cluster-based approach was used to assess differences between TEPs in the pre-drug measurements of the four drug conditions. To adjust for multiple comparisons, a Bonferroni correction was applied to the obtained p-value.

A repeated measure analysis of variance (rmANOVA) with the within-subject effects of DRUG (4 levels: perampanel, dextromethorphan, nimodipine, placebo) and TIME (2 levels: pre-drug, post-drug) was run on the RMT and MEP amplitude data. The Shapiro-Wilk test was applied to test for normal distribution. The MEP data were log-transformed to achieve normal distribution. Sphericity was checked using Mauchly’s test and, whenever violated, the Greenhouse-Geisser correction of the degrees of freedom was applied. For all tests, the significance level was set to p < 0.05.

## Results

TMS was well tolerated by all subjects. In one case, a dosage of 12 mg perampanel caused dizziness, nausea and ataxia, which led to reduction of the dosage to 6 mg for the remaining 13 subjects (i.e., 3 of the reported subjects received 12 mg, the other 13 subjects received 6 mg of perampanel). Otherwise, drugs were well tolerated by all subjects, apart from minor nausea and slight dizziness reported after perampanel and dextromethorphan intake.

### Drug effects on RMT and MEP amplitude

The rmANOVA on RMT values revealed a significant DRUG*TIME interaction (F_3,45_ = 8.993, p < 0.001). *Post hoc* paired t-tests demonstrated a mean RMT increase (post-drug/pre-drug) ± SD after perampanel (1.09 ± 0.08; t_15_ = 4.11, p < 0.001) and nimodipine (1.04 ± 0.04; t_15_ = 2.91, p = 0.007) but not dextromethorphan (0.99 ± 0.07, t_15_ = 0.94, p = 0.36), with compared to RMT change under placebo (0.97 ± 0.07). Importantly, the rmANOVA did not reveal any significant effects of DRUG, TIME or interaction DRUG*TIME on MEP amplitude (Table 1).

**Table 1.** Pre-drug vs. post-drug measurements of RMT (percent maximum stimulator output, %MSO) and MEP amplitudes (mV) (all data, mean ± SD)

| Drug | RMT pre-drug (%MSO) | RMT post-drug (%MSO) | MEP pre-drug (mV) | MEP post-drug (mV) |
| --- | --- | --- | --- | --- |
| Perampanel | 40.0 ± 6.4 | 43.6 ± 7.6 | 0.19 ± 0.31 | 0.16 ± 0.16 |
| Dextromethorphan | 40.9 ± 7.5 | 40.4 ± 6.7 | 0.17 ± 0.35 | 0.18 ± 0.20 |
| Nimodipine | 40.4 ± 7.2 | 41.8 ± 7.1 | 0.18 ± 0.07 | 0.19 ± 0.37 |
| Placebo | 40.8 ± 5.8 | 38.7 ± 8.3 | 0.18 ± 0.21 | 0.16 ± 0.16 |

### TEPs

Pre-drug TEPs and their topographical distributions (**Fig. 2**) were consistent with previous studies of single pulse TMS over M1 (Komssi et al. 2004b; Bonato et al. 2006; Premoli et al. 2014; Darmani et al. 2016). Pre-drug TEPs did not differ between the four drug conditions (all pairwise comparisons, p > 0.05).

**Figure 2.**
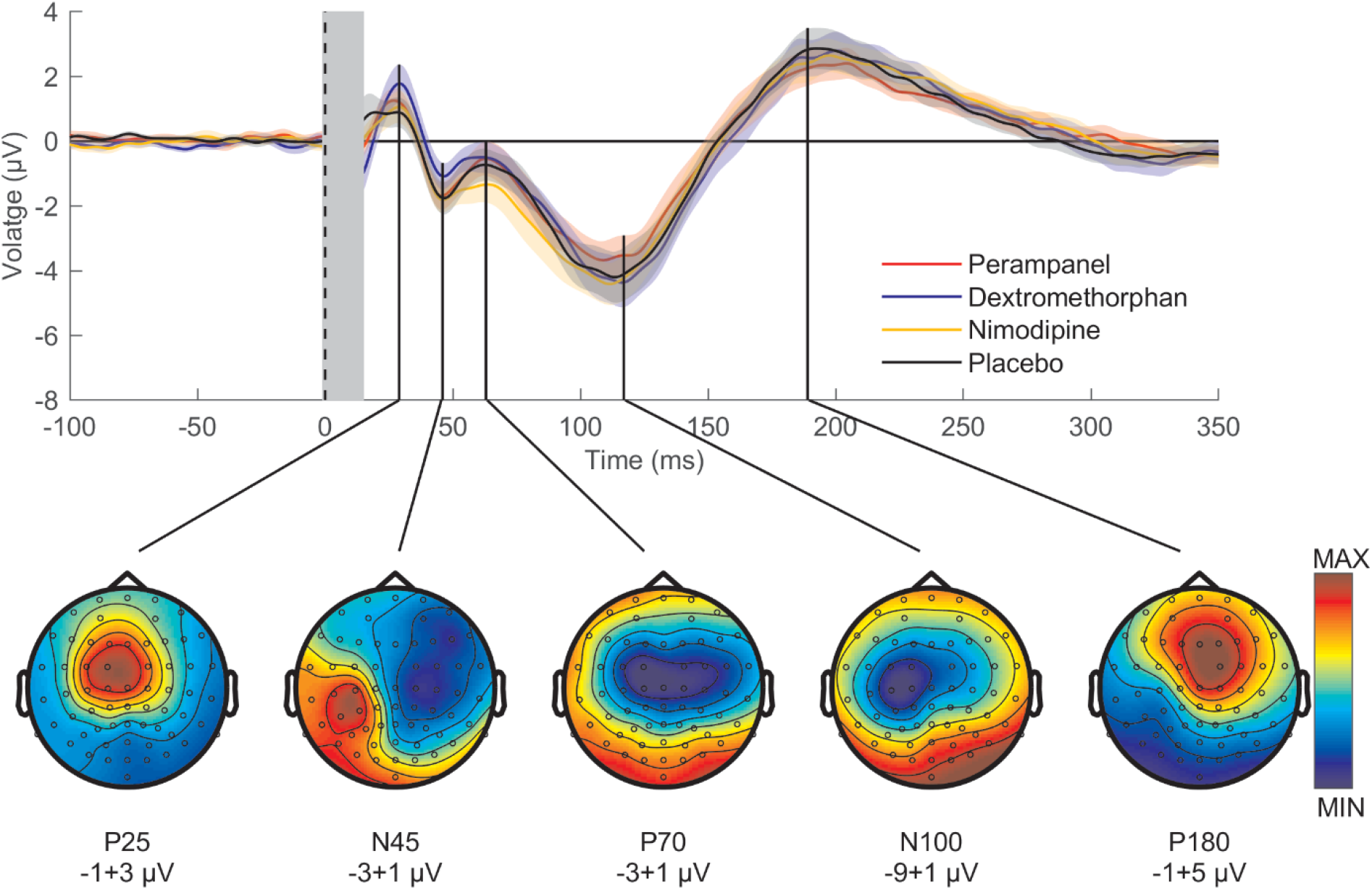
Group average of TEPs evoked by single-pulse TMS of M1 before drug intake. Top panel: pre-drug TEPs averaged across all subjects (n = 16) and EEG electrodes for perampanel (red curve), dextromethorphan (blue curve), nimodipine (yellow curve) and placebo (black curve). Shades represent ±1 SEM. The vertical gray bar represents the time window affected by the TMS artefact that has been removed and interpolated. Bottom panel: pre-drug TEP topographies averaged across subjects (n = 16) and conditions. Each topography was obtained by averaging the signal in the respective TOI (P25: 16-34 ms, N45: 38-55 ms, P70: 56-82 ms, N100: 89-133 ms, P180: 173-262 ms). Data are voltages at sensor level (ranges indicated underneath the plots), while colors are normalized to maximum/minimum voltage.

In the placebo and nimodipine conditions there was no significant difference in the post-drug vs. pre-drug measurement in any of the five TOIs (all p > 0.05; **Fig. 3**). Perampanel resulted in a decrease of the P70 amplitude (p = 0.002; **Fig. 3A, B**). This difference was expressed in predominantly in EEG channels in the non-stimulated hemisphere (**Fig. 4B**, top row). Dextromethorphan increased the N45 amplitude (p = 0.027; **Fig. 3**). The difference was expressed in a bilateral pericentral cluster of electrodes in the stimulated and non-stimulated hemisphere (**Fig. 4B**, bottom row). Single subject data of drug-induced modulations of the P70 and N45 amplitudes are displayed in **Fig. 5** to demonstrate consistency across subjects.

**Figure 3.**
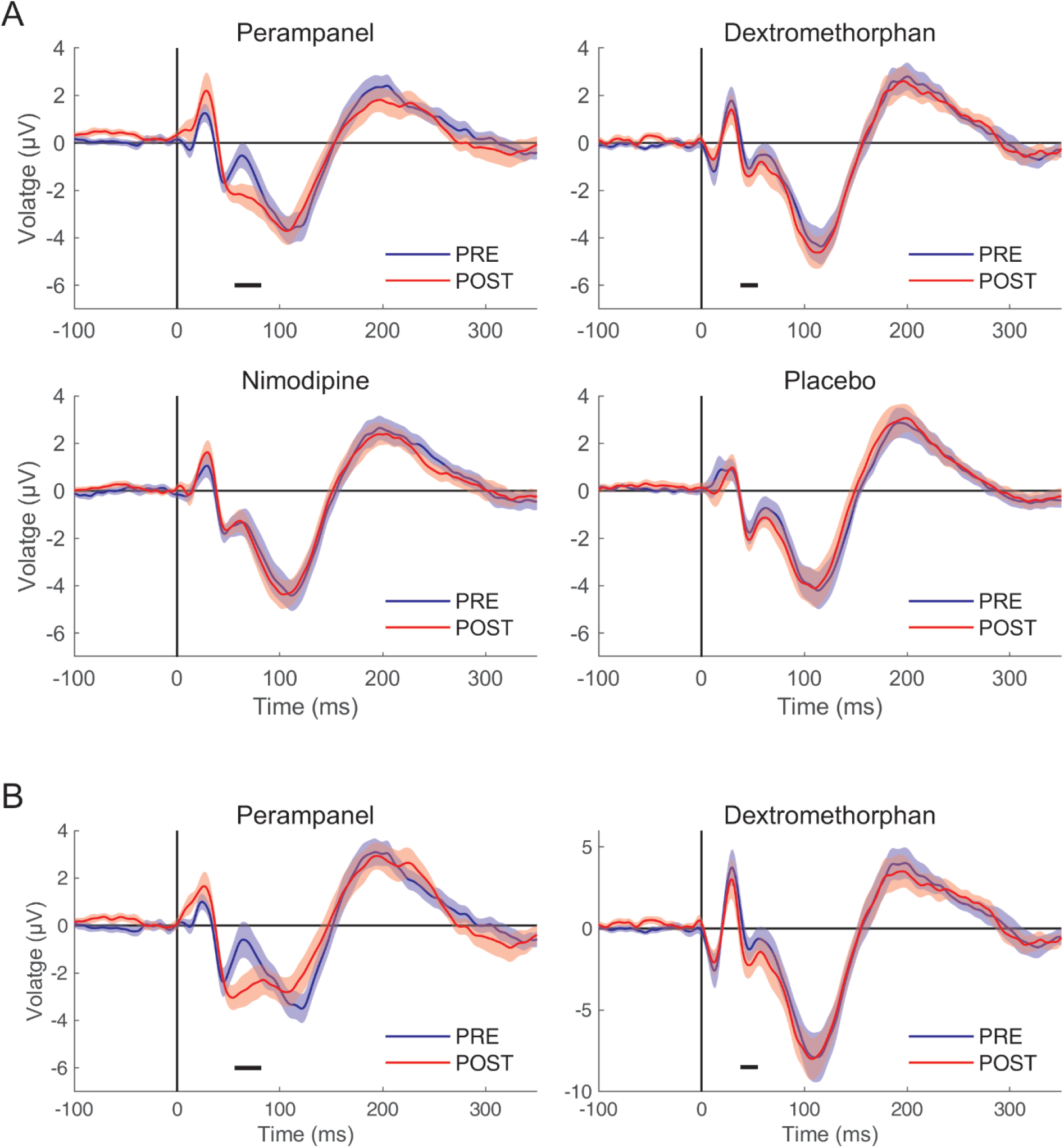
Group average of TEPs pre- and post-drug intake. **(A)** Each panel shows the average TEP time course across subjects and all EEG channels of pre-drug (blue curve) vs. post-drug measurements (red curve) for the four drug conditions. Shades represent ±1 SEM. Significant differences between the pre- and post-drug measurements are indicated with horizontal black bars. **(B)** To better elucidate the drug-induced changes of TEP components shown in (A), the same average TEP time courses are displayed for significant channels only (cf. Fig. 4). Shades represent ±1 SEM.

**Figure 4.**
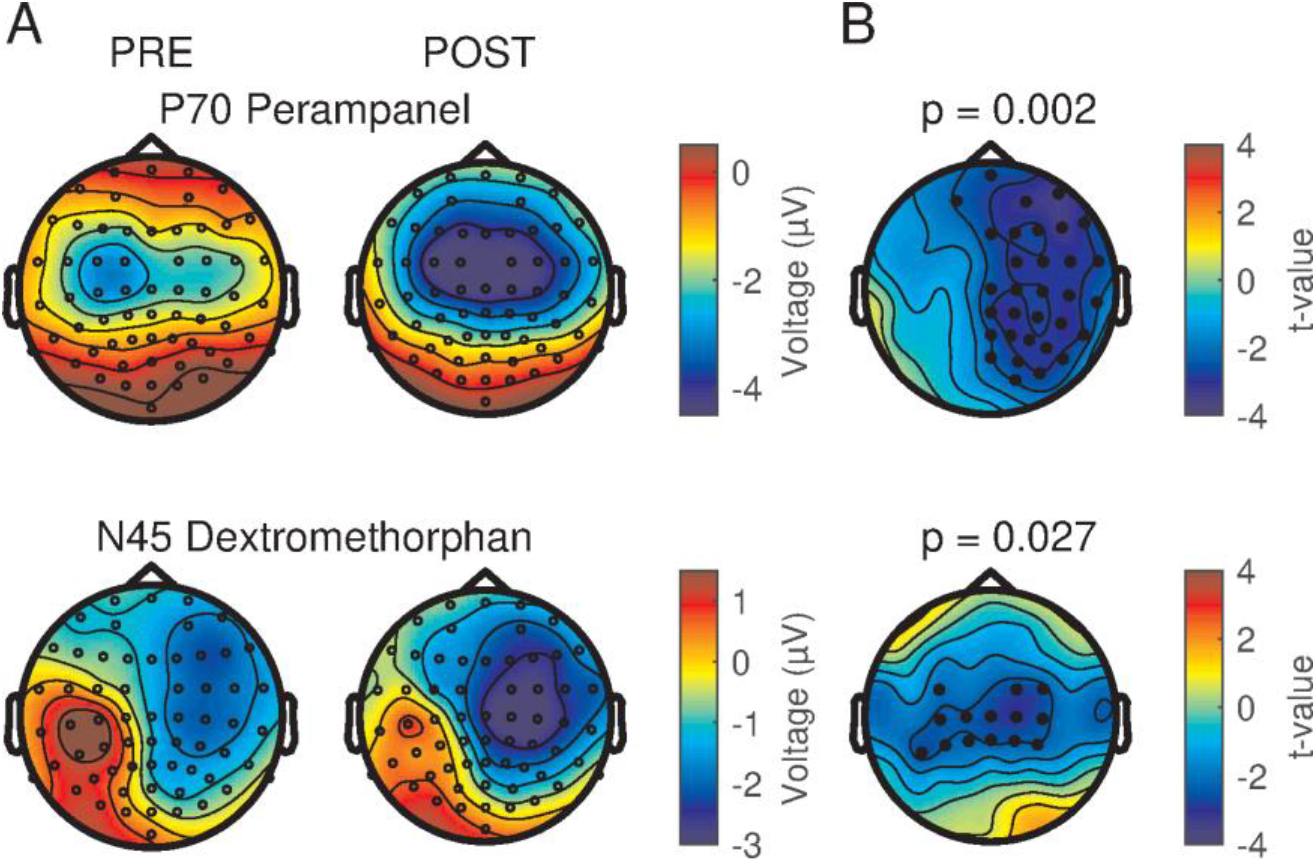
Topographical surface voltage maps for significantly different TEP components. **(A)** Topography of P70 before (left) and after (right) intake of perampanel (top row) and topography of N45 before (left) and after (right) intake of dextromethorphan (bottom row). **(B)** T-value statistical maps with channels belonging to significant clusters highlighted as black dots.

**Figure 5.**
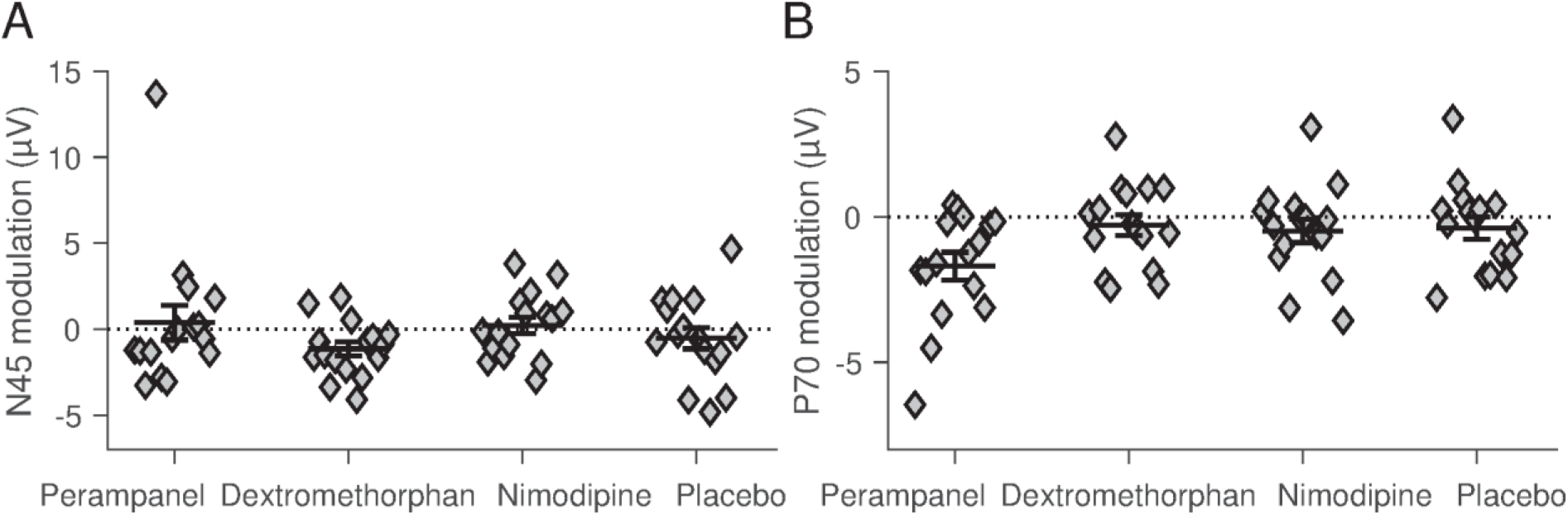
Scatter plots of single subject drug-induced TEP changes. Amplitude modulations (post-drug minus pre-drug) of the N45 (***A***) and P70 (***B***) TEP components for perampanel, dextromethorphan, nimodipine and placebo. For the investigated TEP components, amplitudes were calculated as the average voltage for identified significant channels for dextromethorphan (N45) and perampanel (P70). Error bars indicate mean ±1 SEM.

## Discussion

In this study, we investigated modulation of TMS-evoked EEG potentials by a single oral dose of an AMPA receptor antagonist (perampanel), an NMDA receptor antagonist (dextromethorphan), and an L-VGCC blocker (nimodipine). Perampanel decreased the P70 amplitude, whereas dextromethorphan increased the N45 amplitude. Nimodipine and placebo had no effect on TEP amplitudes. Our results show specific modulation resulting from drugs that act on glutamate receptors. The differential effects are likely caused by differences in the specific modes of drug action, as discussed in detail below.

### N45 modulation by dextromethorphan

Dextromethorphan binds preferentially to NMDA receptors and reduces EPSPs due to inhibition of calcium influx. In paired-pulse TMS-EMG studies, dextromethorphan decreased intracortical facilitation (Ziemann et al. 1998), a marker of glutamatergic neurotransmission (Ziemann et al. 2015), while it did not affect RMT or MEP amplitude (Ziemann et al. 1998; Fitzgerald et al. 2005; Wankerl et al. 2010). This pattern of effects on TMS-EMG measures is like the one of benzodiazepines, which also reduce intracortical facilitation (Ziemann et al. 1996; Ziemann et al. 2015), probably through enhancement of short-interval intracortical inhibition, a marker of GABAAergic inhibitory postsynaptic potentials (IPSPs) that superimposes with intracortical facilitation (Hanajima et al. 1998). At the level of TMS-EEG measurements, dextromethorphan showed a virtually identical effect as benzodiazepines (Premoli et al. 2014; Premoli et al. 2018) by increasing the N45 amplitude (cf. Fig. 6). Therefore, the present data lead to the proposition that the N45 amplitude reflects excitation-inhibition balance of EPSPs and IPSPs evoked by the TMS pulse. This significantly extends the previous view that the N45 amplitude exclusively reflects GABAAergic inhibition (Premoli et al. 2014; Darmani et al. 2016; Premoli et al. 2018).

**Figure 6.**
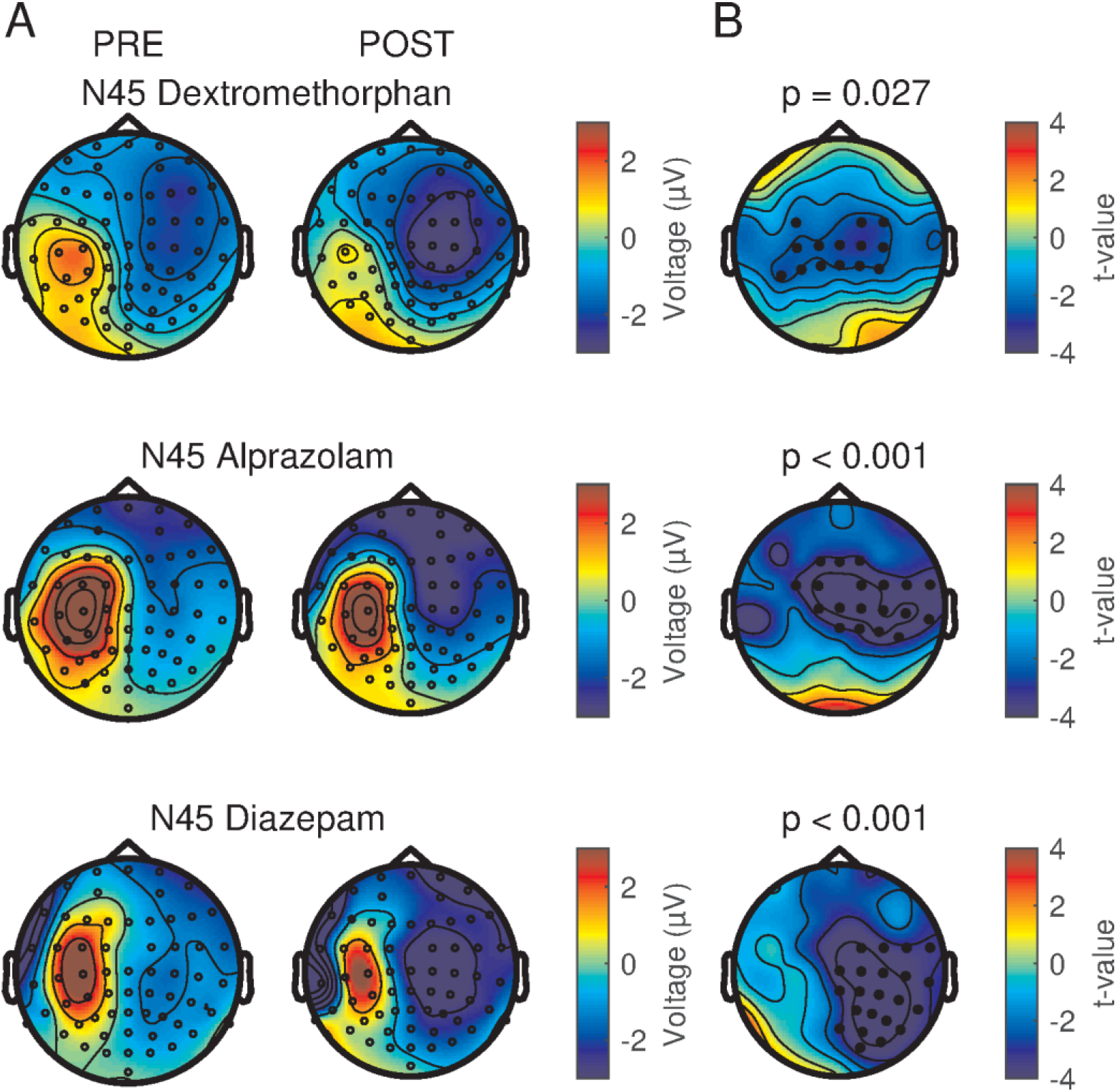
Comparison of the modulation of the N45 TEP by dextromethorphan and two classical benzodiazepines (alprazolam and diazepam, results adapted from (Premoli et al. 2014)). First two columns show voltage surface maps of N45 recorded before and after drug intake. The third column shows t-statistic maps of the N45 cluster post-drug versus pre-drug differences. Electrodes of the significant clusters are denoted by black dots.

Of note, while the enhancing effects of the NMDA receptor antagonist dextromethorphan and benzodiazepines on the N45 amplitude are similar, dextromethorphan (and perampanel) had no effect on the N100 amplitude in the non-stimulated hemisphere, while benzodiazepines decreased it (Premoli et al. 2014; Premoli et al. 2018). Together, these findings support the idea that the N100 in the frontal cortex of the non-stimulated hemisphere reflects propagated neural activity controlled by GABAAergic but not glutamatergic neurotransmission.

### P70 modulation by perampanel

AMPA receptor activation in response to glutamate binding generates fast EPSPs followed by rapid current decay (Niciu et al. 2012). The effect of the AMPA receptor antagonist perampanel was specific by reducing the P70 amplitude. Importantly, this effect was almost exclusively expressed in the non-stimulated right hemisphere (cf. Fig. 4), suggesting that the effect of perampanel is specific on interhemispherically propagated neural activity. This finding is in close agreement with intrahemispheric and interhemispheric spread of epileptiform activity in rodent cortical slices that was not influenced by application of the NMDA receptor antagonist D-2-amino-5-phosphonovaleric acid (D-APV), but blocked by the AMPA receptor antagonist 6-cyano-7-nitroquinoxaline-2,3-dione (CNQX) (Alefeld et al. 1998; Telfeian and Connors 1999). The P70 has not shown reactivity to any other of the so far tested drugs (positive allosteric modulators at the GABAA receptor, alpha-5 GABAA receptor antagonist, GABAB receptor agonist, voltage-gated sodium channel blockers, NMDA receptor antagonist, L-VGCC blocker) (Premoli et al. 2014; Darmani et al. 2016; Premoli et al. 2017; Premoli et al. 2018). We therefore propose that the P70 amplitude reflects glutamatergic (interhemispheric) signal propagation mediated by AMPA receptor activation. Whether the P70 amplitude is exaggerated in epilepsy, and may be used as a biomarker to predict antiepileptic drug responses, is currently unclear, as the very few available TEP studies were performed exclusively in patients with generalized epilepsies on antiepileptic drug treatment, without alteration of the P70 amplitude (Julkunen et al. 2013; Ter Braack et al. 2016; Kimiskidis et al. 2017).

### Absence of TEP modulation by nimodipine

L-VGCCs are expressed on dendrites of neurons throughout the central nervous system. They contribute to regulation of neuronal excitability. L-VGCCs open from their closed/resting state only upon strong postsynaptic depolarization (Nowycky et al. 1985). In addition, L-VGCCs are not significantly involved in controlling glutamate release from presynaptic nerve terminals (Catterall 2011). Therefore, L-VGCCs should not play a role in the initial excitation of neurons by the TMS pulse in resting motor cortex. Accordingly, nimodipine had no or only very minor effects on RMT or MEP recruitment curve in the present and previous studies (Wankerl et al. 2010; Weise et al. 2017), and did not show any effect on TEPs in the present study. Finally, a failure to obtain a nimodipine effect on TEPs due to a too low dosage can be largely excluded, as the same single oral dose of 30 mg resulted in significant suppression of long-term potentiation and long-term depression-like plasticity in human motor cortex (Wolters et al. 2003; Wankerl et al. 2010; Weise et al. 2017).

## Conclusions

Findings support the general notion that TEPs evoked by single-pulse TMS of M1 can be used as markers of excitability and propagated neural activity in the human brain. Specifically, the effects of the NMDA receptor antagonist dextromethorphan extend our understanding of the N45 potential to reflect excitation-inhibition balance regulated by NMDA and GABAA receptors. Furthermore, the suppressive effects of perampanel on the P70 potential in the non-stimulated hemisphere support the idea that this propagated activity is controlled by glutamatergic neurotransmission through AMPA receptors. Finally, the null effects of the L-VGCC blocker nimodipine on TEPs are in accord with the known physiology of L-VGCCs on neuronal excitability. Altogether, pharmaco-TMS-EEG advances our knowledge of the physiology underlying TEPs, and this may be of directly utility in interpreting TEP abnormalities in neurological and psychiatric disorders with pathological neural excitability or signal propagation in brain networks.

## Acknowledgements

This work was supported by a grant from the German Research Foundation (DFG ZI 542/9-1) to U.Z.

## Conflicts of Interest

U.Z. received grants from the German Ministry of Education and Research (BMBF), Biogen Idec GmbH, Servier, and Janssen Pharmaceuticals NV, and consulting fees from Biogen Idec GmbH, Bayer Vital GmbH, Bristol Myers Squibb GmbH, Pfizer GmbH, CorTec GmbH, and Medtronic GmbH, all not related to this work. All other authors declare that they have no conflicts of interest.

