## Supplemental Tables1 and 2 for "TMS-EEG signatures of glutamatergic neurotransmission in human cortex"

**Supplementary Table 1.** Drug dosages, way of application (Route), time of peak plasma concentration after intake (T_max_) and half-life time (T_1/2_) of the study drugs.

| **Drug** | **Brand Name** | **Dosage** | **Route** | **T_max_ [h]** | **T_1/2_ [h]** |
| --- | --- | --- | --- | --- | --- |
| Perampanel | Fycompa^®^ | 12 (or 6) mg | tablet | 0.5-4^1^ | 95^1^ |
| Dextromethorphan | Hustenstiller-ratiopharm^®^ Dextromethorphan | 120 mg | capsule | 1-2^2^ | 1.2-2.2 hrs^3^ (CYP2D6-EM)  <45 hrs^3^ (CYP2D6-PM) |
| Nimodipine | Nimodipin-HEXAL^®^ | 30 mg | tablet | 0.6-1.6^4^ | 1.1-1.7^4^ |
| Placebo | P-Tabletten Lichtenstein, 7, 8, 10 mm, Winthrop | n.a. | tablet /  capsule | n.a. | n.a. |

^1^ Phase I randomized biopharmaceutical study E2007-A001-040 conducted by Eisai (personal communication with Eisai), ^2^ PRODUCT INFORMATION Hustenstiller-ratiopharm® Dextromethorphan, ^3^ Vetticaden SJ (1989) Pharm Res 6:13-9, ^4^ PRODUCT INFORMATION NIMOTOP® Nimodipine Bayer Resources. n.a., not applicable.

**Supplementary Table 2.** Drug administration

| **Drug condition** | **0 min after pre-drug**  **measurements** | **60 min after pre-drug measurements** |
| --- | --- | --- |
| Perampanel | 4 capsules Placebo | 1 tablet Fycompa® 12 (or 6) mg |
| Dextromethorphan | 4 capsules Hustenstiller-ratiopharm® Dextromethorphan 30 mg | 1 tablet Placebo |
| Nimodipine | 4 capsules Placebo | 1 tablet Nimodipin-HEXAL® 30 mg |
| Placebo | 4 capsules Placebo | 1 tablet Placebo |

Study drugs were given orally in randomized order on separate study visits spaced by at least two weeks according to the above scheme (at each study visit, 4 capsules directly and 1 tablet 60 min after the end of the pre-drug measurements, respectively) to ensure double-blinded state for subjects and investigators.
